# Genetic and thermal variation influence adaptation to fluctuating temperature in the seed beetle, *Callosobruchus maculatus*

**DOI:** 10.1101/2021.11.26.470113

**Authors:** Edward Ivimey-Cook, Claudio Piani, Wei-Tse Hung, Elena C. Berg

## Abstract

Climate change is associated with both the increase in mean and variability of thermal conditions. The use of more realistic thermal regimes is therefore the most appropriate laboratory method to correctly predict population responses to thermal heterogeneity. However, both the long- and short-term implications of evolving under such conditions are not well understood. Here, we examined the effect of fluctuating daily temperatures on several key life history traits in the seed beetle, *Callosobruchus maculatus*, that was exposed to a short-term thermal switch into a novel environment. Populations were kept for 19 generations at one of two temperatures: constant control temperature (T=29°C) or a fluctuating daily cycle (T_mean_=33°C, T_max_=40°C, and T_min_=26°C) and were subsequently exposed to a switch to the opposite condition. We found that beetles that had evolved in stressful environments were smaller in size when switched to a constant 29°C and had far greater reproductive fitness compared to beetles from both the constant control and continuously stressful 33°C environments. This suggests that beetles raised in environments with stressful fluctuating temperatures were more phenotypically plastic and had greater genetic variability than control treatment beetles and indicates that populations that experience fluctuations in temperature may be better able to respond to short-term changes in environmental conditions.

## Introduction

A multitude of studies have shown the negative impact of increasing mean temperature on important life history traits in invertebrate species (Bauerfeind & Fischer, 2014; Rogell *et al.*, 2014; Vasudeva *et al.*, 2014; Berger *et al.*, 2017; Chen *et al.*, 2018; Klepsatel *et al.*, 2019; Ivimey-Cook *et al.*, 2021). In particular, heat stress caused by high temperatures outside of a species’ thermal optimum can disrupt reproduction, reduce longevity and survival in insects (Berger *et al*., 2017; Ivimey-Cook *et al.*, 2021), particularly if outside of a species Thermal Fertility Limit (TFL, reviewed in Walsh *et al.*, 2019).

Increasing mean temperatures, as well as the increasing frequency of Extreme High Temperature events, (EHTs, Ma *et al.*, 2021), are two of the chief outcomes of rising anthropogenic gas emissions that characterise global climate change (Lashof & Ahuja, 1990; Pereira *et al.*, 2012; Pachauri *et al.*, 2014; Ummenhofer & Meehl, 2017). This change in temperature and frequency of EHTs can influence populations both directly, for example, by increasing the rate of genome wide *de novo* mutations (Berger *et al.*, 2017, 2021), or indirectly, for example, by triggering phenological mismatches between species and prey (Visser *et al.*, 1998, 2012; see also Parmesan, 2006 for an extensive review). This increase in environmental stress leads populations to either adapt and track environmental conditions through phenotypic plasticity, to shift their geographic boundaries to match thermal tolerance (either by changing latitude or altitude), or to go extinct (Holt, 1990; Pereira *et al.*, 2012; Hetem *et al.*, 2014; Kellermann *et al.*, 2019; Lehmann *et al.*, 2020; Trisos *et al.*, 2020). Until recently, species in tropical habitats were predicted to be especially vulnerable, as they were thought to have evolved far more narrow and specialized thermal ranges (Janzen, 1967; Deutsch *et al.*, 2008; Walters *et al.*, 2012). However recent work using multiple *Drosophila* species (n = 22) has questioned this idea, with results showing that the breadth of thermal performance and point of thermal optimum did not differ between temperate and tropical species (MacLean *et al.*, 2019).

Not only is the Earth’s mean temperatures rising, but it is also becoming more variable (Folguera *et al.*, 2011; Vasseur *et al.*, 2014; Masson-Delmotte *et al.*, 2021). This increase in environmental variability is likely to intensify the already detrimental impacts of elevated temperatures (Vasseur *et al.*, 2014). Building on work by Deutsch *et al.* (2008), Vasseur *et al.* (2014) found that changes to both the mean and variance in environmental temperature produced long-term changes to population fitness. More specifically, performance data collected from 38 globally distributed invertebrate species indicated that whilst increasing mean temperature had a mixture of both positive and negative effects on performance, which were largely latitude-dependent, increased variation was associated with more negative effects (Vasseur *et al.*, 2014). In addition, the authors highlighted the relative contribution of mean and variance in determining projected performance in the years 2050-2059. They found that a change in mean temperature explained only 32% of future species performance, whereas combining both the mean and variance simultaneously explained 93%. This highlights the importance of understanding how species react not only to changes in mean temperature, but also to changes in temperature *variation* in order to form accurate predictions of a population’s response to climate change.

Until recently, studying the effects of climate warming and the resulting adaptive responses of organisms was confined to field studies, with little control over multiple variables, or laboratory studies that maintained populations at constant temperatures (Fischer *et al.*, 2011; Folguera *et al.*, 2011; Thompson *et al.*, 2013; Bauerfeind & Fischer, 2014). However, we now know that populations often exhibit marked life history differences when exposed to fluctuating rather than constant thermal stress in the laboratory (Folguera *et al.*, 2011; Bauerfeind & Fischer, 2014; Kellermann & van Heerwaarden, 2019; Buckley & Kingsolver, 2021). For example, reaction norms of the mosquito *Anopheles stephensi* under fluctuating thermal regimes were fundamentally different to those under constant temperatures (Paajimans *et al.*, 2013). Similarly, in the tropical butterfly *Bicyclus anyana*, life history traits were strongly influenced by both the mean and the amplitude of the experimental thermoperiod (Brakefield & Mazzotta, 1995). Indeed, the use of constant temperatures might significantly underestimate the impact of global warming on life history evolution (Fischer *et al.* 2011). Fortunately, there is now a well-accepted requirement that empirical studies incorporate more realistic thermal regimes into their experimental design (Niehaus *et al.*, 2012; Paaijmans *et al.*, 2013; Buckley & Kingsolver, 2021). As a result, the number of laboratory studies involving invertebrates and “natural” thermal variation is ever increasing (Colinet *et al.*, 2015; Morash *et al.*, 2018; Ma *et al.*, 2021b; Morgan Fleming *et al.*, 2021; Chang *et al.*, 2022). However, there is still the need to understand the long-term implications of evolving under natural conditions that mimic the thermal stress associated with climate warming on the short-term ability to adapt to novel temperatures, and contrasting these outcomes with populations evolving under constant stress-free conditions.

To this end, we investigated the long-term effects of thermal stress caused by realistically fluctuating temperatures on the phenotypic plasticity and adaptation of two laboratory populations of seed beetle, *Callosobruchus maculatus.* In addition, we examined the short-term effects of exposure to a novel environment both into and from the fluctuating environment. Two different thermal regimes were used: in one regime, beetles were kept at a constant control temperature of 29°C, and in the second regime we exposed beetles to a more stressful and realistic fluctuating daily cycle with a mean temperature of 33°C (representative of a May day in southern India, where the laboratory lines were originally collected). The two beetle populations were then either kept in these two environments for the entirety of the experiment or were *switched* to the other environment for two generations without selection prior to assaying several key life history traits (development time, body mass and various measures of reproductive fitness). Previous work in the same species of beetle has highlighted the negative effect of increasing temperature with environmental fluctuation on fecundity traits (with little to no effect on hatching success, development time and body mass; Hallsson & Björklund, 2012). Importantly, our definition of fluctuating environments differs from theirs. Whilst the term employed here represents daily sinusoidal fluctuations around a constant mean, representative of the natural environment that a seed beetle could experience, Hallsson and Björklund (2012) used the term to describe the addition of thermal noise around a linear increase from 30°C to 36°C. Thus, we predict that beetles will exhibit entirely different life-history responses when populations are exposed to stressful but repeating sinusoidal fluctuations without a corresponding linear increase in mean temperature.

## Methods

### Study system

*C. maculatus* is an agricultural pest that originates from Africa and Asia but is currently found throughout the world. Adult seed beetles are facultatively aphagous – that is, they do not require food or water to survive and acquire all the resources they need during the larval stage within the bean. Aphagous adults can live up to two weeks but adults with access to nutrients can live three weeks or more (Fox, 1993; Ursprung *et al.*, 2009). Females lay their eggs on the surface of dried legumes, such as mung beans (*Vigna radiata*) or black-eyed beans (*Vigna unguiculata*). Larvae from the fertilized eggs burrow into the bean and eclose as adults 23-27 days later. Females of this species start to mate and lay eggs as soon as they emerge (Beck & Blumer, 2014).

### Laboratory evolution lines and climatic conditions

In our experiment, we used two different strains of *C. maculatus*, South India (SI) USA and SI Leicester, both derived from a population of beetles collected in Tirunelveli, India, in 1979 (Mitchell, 1991) and then subsequently raised at the University of Kentucky, USA, and the University of Leicester, UK (from 1992), respectively (Fricke & Arnqvist, 2004). We obtained these strains from Uppsala University in Sweden and maintained them at the American University of Paris for approximately 25 generations before establishing the replicates for the current experiment. Beetles were cultured exclusively on mung beans and kept in climate chambers at a constant 29°C, 50% relative humidity, and a 12:12h light:dark cycle. They were housed at aphagy in 1L jars with 250g of beans, and approximately 250-350 newly hatched beetles were transferred to new jars with fresh beans every 24 days on a continual basis.

For 19 generations prior to the experiment, we subdivided SI USA and SI Leicester populations (Fig. S1A) into four replicates each. Two “Control” replicates of each were maintained at a constant temperature of 29°C, while two “Stress” replicates were subject to a daily temperature cycle (Fig. S1B) consisting of 12 separate 2hr periods of constant temperature *T_i_*,

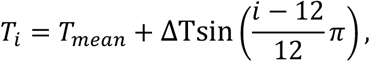

where *T_mean_* = 33 °C, Δ*T* = 7°C, and i = 0,1…. 11. This is a stepwise sinusoidal temperature cycle with T_max_=40 and T_min_=26 that mimics typical late spring condition in Southern India, where this species evolved. Humidity and light cycles remained the same for all replicates: 50% relative humidity, and a 12:12 h light:dark cycle.

Prior to the assays of body size, development time, and reproductive fitness (see next section), we created two replicates of two populations (SI USA and SI Leicester; see Fig. S1A/B) for:

1. beetles adapted to and assayed at Control conditions (**USA**-C and **LEI**-C);
2. beetles adapted (for one year) to and assayed in Stress conditions (**USA**-S and **LEI**-S);
3. beetles adapted to Control conditions that were then acclimatized to Stress conditions for two generations without selection prior to being assayed (**USA**-C-to-S and **LEI**-C-to-S); and
4. beetles adapted to Stress conditions that were acclimatized to Control conditions for two generations without selection prior to being assayed (**USA**-S-to-C, **LEI**-S-to-C).

To separate genetic adaptation from phenotypic plasticity – that is, to control for the influence of parental effects – we created “acclimatized” treatments by allowing beetles from one set of conditions to adjust to the new set of conditions for two generations prior to assays (C-to-S and S-to-C; Fig. S1B). Critically, to remove the possible effects of natural selection and rule out the possibility that rapid genetic responses might influence our measurements for the “acclimatized” treatments, we randomly paired 50 pairs of virgin beetles during each of the two generations. That is, we removed the effects of assortative mating.

For the Stress-to-Control (USA S-to-C, LEI S-to-C) and the Control-to-Stress (USA C-to-S, LEI C-to-S) treatments, the following steps were performed. First, we moved beans with fertilized eggs from jars into 48-well virgin chambers. When enough males and females had hatched (from day 21-24 after hatch), we paired 60 males and 60 females in 60 60-mm petri dishes with 100 beans in each dish and placed these dishes in the opposing chamber. We made sure that there were sufficient number of beans in each dish for females to lay just one egg per bean, thereby eliminating competition among multiple larvae within a bean. To further reduce larval competition, we removed adult beetle pairs from the petri dishes after 72 hours.

After 19 days, we then chose the 50 petri dishes with the most fertilized eggs and put 48 fertilized eggs from each petri dish into individual marked 48-well virgin chambers. One day after eclosion, we randomly selected and paired 50 females and 50 males and put them into 50mm petri dishes with 80 beans. After 72 hours, the adult males and females were removed from the petri dishes. After 19 days, beans with fertilized eggs from each petri dish were transferred into 48-well virgin chambers. We then marked the date of hatch and sex of all offspring and conducted assays of development time, body mass, and reproductive fitness.

For the continuous C and S treatments, we skipped the acclimatization steps and simply moved beans with fertilized eggs into virgin chambers prior to hatch, selecting one-day old males and females for all subsequent assays.

### Body mass assay

Across two replicates, 8 males and 8 females were weighed for each of the 16 lines (2 populations, 2 sexes, 4 temperature treatments; note for USA-C, 16 males and 16 females were weighed) using an Ohaus Pioneer Plus Analytical Balance (Model PA214C). All virgin chambers were weighed empty, prior to adding the beetles to be weighed. We then calculated the difference between the weight of the virgin chamber before the beetles were placed inside and the weight of the virgin chamber after the beetles were placed inside. Beetles from the body mass assay were not used for the other assays.

### Development time and fitness assays

Development time and fitness assays were done concurrently. We paired 50 1-day old male and female beetles and placed them into 60mm petri dishes filled with 85 beans. After 24 hours, each pair was moved together into a new 60mm petri dish filled with 75 beans, and the first dish was set aside in the same climate chamber. The next day, this procedure was repeated. The beetles were left in the third and final set of dishes until they died. The first dishes for each pair were marked “Day 0”, the second dishes were marked “Day 1”, and the third and final dishes were marked “Day 2+”. After 19 days for each dish, all the beans with fertilized eggs were placed into virgin chambers, which were monitored daily for the date of eclosion and the sex of all offspring. To calculate development time, we counted the number of days between the date that an egg was laid and the date the adult offspring eclosed. To calculate LRS, we calculated the number of male and female offspring that emerged for each pair, from all three sets of dishes (Days 0, 1, and 2+). To calculate individual fitness (or lind), we took the dominant eigenvalue (using the popbio v. 2.7 package, (Stubben & Milligan, 2007) from an age-structured Leslie matrix (Leslie, 1945) where the top row denotes fertility and the sub-diagonal represents survival from age *t* to *t*+1. 24 days of development time were added prior to day 0-2+ fertility schedule.

### Statistical analyses

All statistical analyses were performed in R v. 4.1.0 (R Core Team, 2016) using glmmTMB v. 1.0.2.9000 (Brooks *et al.*, 2017; Magnusson *et al.*, 2019) with pairwise interactions and slopes compared using emmeans v. 1.6.1 (Lenth *et al.*, 2019). Ggplot2 v. 3.3.4 (Wickham, 2009) was used for all graphical visualization. The overall effects of each variable were identified using the Type 3 Anova function from car v. 3.0-10 package (Fox *et al.*, 2012).

For development time, body mass, and individual fitness, a Gaussian linear mixed effects model was fitted (after log-transforming development time to reduce right-skew) with the fixed effects of Population, Treatment, Sex (for development time and body mass), and all higher order interactions. In addition to these main effects, a fixed two-level factor of replicate number was added. Lastly, for the models involving development time, the random effect of parental ID was added to account for potential pseudoreplication. If the three-way interaction was non-significant (identified from the Anova function), it was removed in order to reduce model complexity. A generalized linear mixed effects model with Poisson distribution and similar fixed effect structure was fitted for LRS and age-specific reproduction, albeit without the variable Sex and all subsequent two/three-way interactions. In addition to a covariate of Day (and all higher-order interactions with Population and Treatment), a random effect of individual ID was added to the age-specific reproduction model to account for repeated measures. Both models were checked for overdispersion and zero-inflation (ZI) using DHARMa v. 0.4.1, (Hartig, 2020) by simulating the residuals of a Poisson model. If significant, a variety of error distributions with/without zero-inflated parameters (if ZI was identified) were then fitted and the best model identified by Akaike’s Information Criterion (AIC).

## Results

### Body mass

Individual body mass was significantly affected by both of the interactions involving Sex (Sex and Population: χ^2^ (1) = 25.95, p<0.001; Sex and Treatment: χ^2^(3) = 19.18, p<0.001; Table S1A) but not by the remaining two-way interaction of Population and Treatment (χ^2^(3) = 2.51, p = 0.47) or the three-way interaction (χ^2^ (3) = 6.37, p = 0.09). Females were generally heavier than males (Fig. 1) in both populations. The difference was more pronounced in the USA population (Average body mass (g): F-LEI = 0.087, M-LEI = 0.075, estimate = 0.013, p<0.001; F-USA = 0.089, M-USA = 0.068, estimate = 0.021, p<0.001; Fig. 1, Tables S1A-B). The females of both populations were similar in weight, but the USA males were smaller (F-LEI = 0.087, F-USA = 0.089, estimate = −0.002, *p* = 0.35; M-LEI = 0.075, M-USA = 0.068, estimate = 0.007, p<0.001; Fig. 1, Tables S1A-B). The relative difference between male and female weight was largely dependent on temperature treatment (Fig. 1). Overall, beetles raised in the S-to-C regime were smaller for both sexes in comparison to other treatments (all comparisons significant to p = 0.05; Tables S1A-B). However, the difference between male and female weights was far less pronounced in the constant stress (S) treatment (albeit all differences were still significant to p<0.001; F-S-to-C = 0.083, F-S = 0.089, F-C-to-S = 0.091, F-C = 0.089; M-S-to-C = 0.064, M-S = 0.079, M-C-to-S = 0.071, M-C = 0.071; Tables S1A-B).

**Figure 1.**
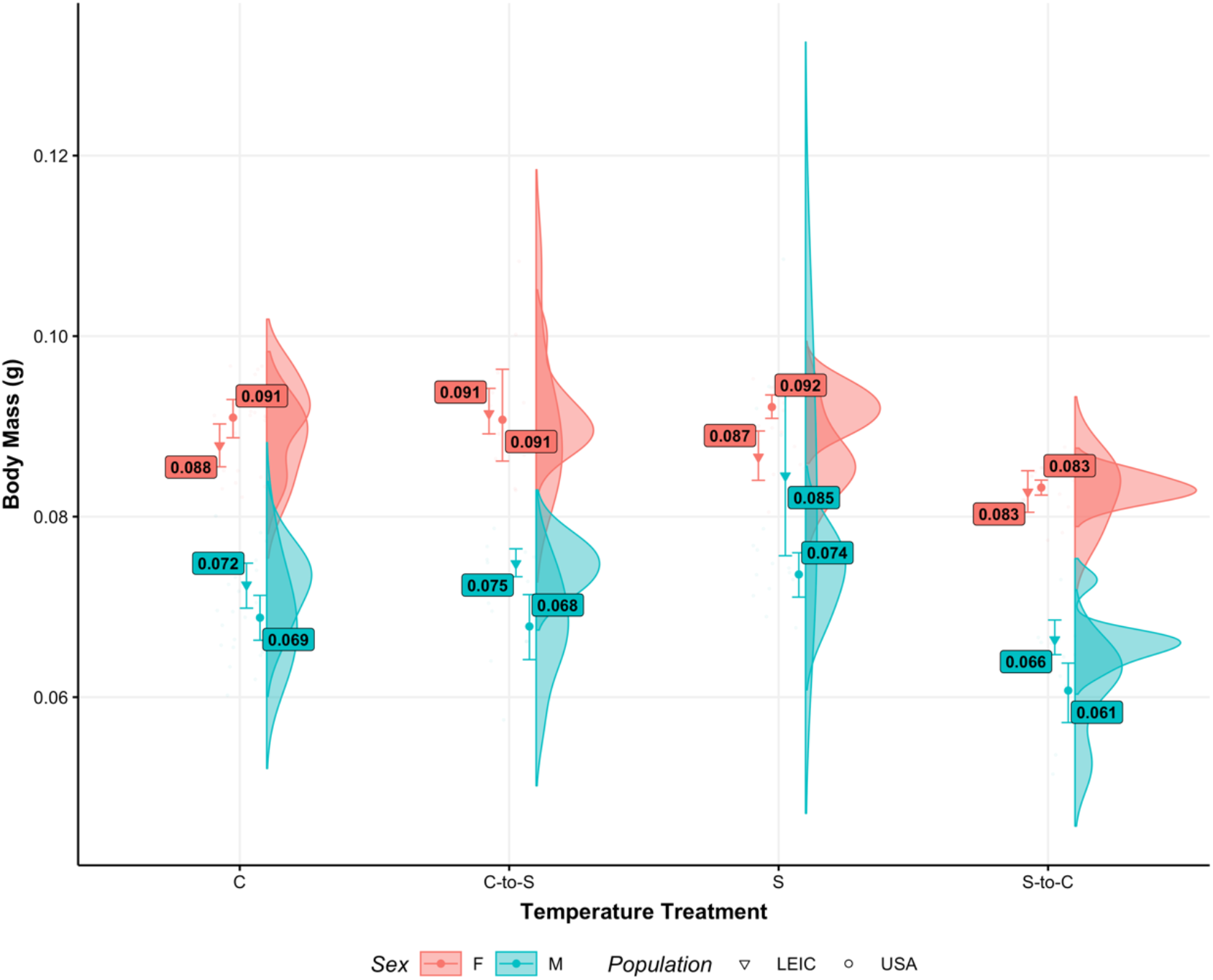
Body mass of individuals from the Leicester (triangle) or USA (circle) populations in the C (Control), C-to-S (Control-to-Stress), S (Stress), or S-to-C (Stress-to-Control) temperature treatments. Sex is denoted with either blue (male) or red (female) points. Points represent mean values with accompanying standard errors. Half violins show the distribution of raw data.

### Development time

Development time was influenced by both two-way interactions involving Temperature treatment (Treatment and Population: χ^2^ (3) = 105.94, *p* <0.001; Treatment and Sex: χ^2^ (3) = 11.76, *p* = 0.008). Similar to body mass, no detectable three-way interaction was found (χ^2^ (3) = 1.71, p = 0.635). The USA and Leicester populations responded differently to the four temperature treatments, with beetles from the Leicester population developing faster on average. Populations from USA took significantly longer to develop in comparison to populations from Leicester in both the Control and S-to-C Treatments (Average Development Time (Days): LEI-C = 23.7, USA-C = 24.7, Ratio = 0.960, *p*<0.001; LEI-S-to-C = 23.1, USA-S-to-C = 24.1, Ratio = 0.961, *p*<0.001; Fig. 2) but not in C-to-S (LEI-C-to-S = 23.0, USA-C-to-S = 23.0, Ratio = 0.999, *p* = 1.00) or S Treatments (LEI-S = 22.5, USA-S = 22.3, Ratio = 1.01, *p* = 0.38; Fig. 2, Tables S2A-B). Despite these differences, both populations in the S treatment developed faster compared to other temperature treatments (LEI-S = 22.5, LEI-S-to-C = 23.1, LEI-C-to-S = 23.0, LEI-C = 23.7; USA-S = 22.3, USA-S-to-C = 24.1, USA-C-to-S = 23.0, USA-C = 24.7). In addition, whilst males typically developed faster in both populations (shown with lower mean values for 6 out of 8 population/treatment combinations, Fig. 2), only in the S-to-C treatment was this difference significant (F-S-to-C = 23.7, M-S-to-C = 23.50, Ratio = 1.008, *p*<0.001; Fig. 2, Tables S2A-B). However, both males and females in the S treatment developed faster than other temperature treatments and showed no sex difference in development time (F-S = 22.4, F-S-to-C = 23.7, F-C-to-S = 23.0, F-C = 24.2; M-S = 22.4, M-S-to-C = 23.5, M-C-to-S = 22.9, M-C = 24.1).

**Figure 2.**
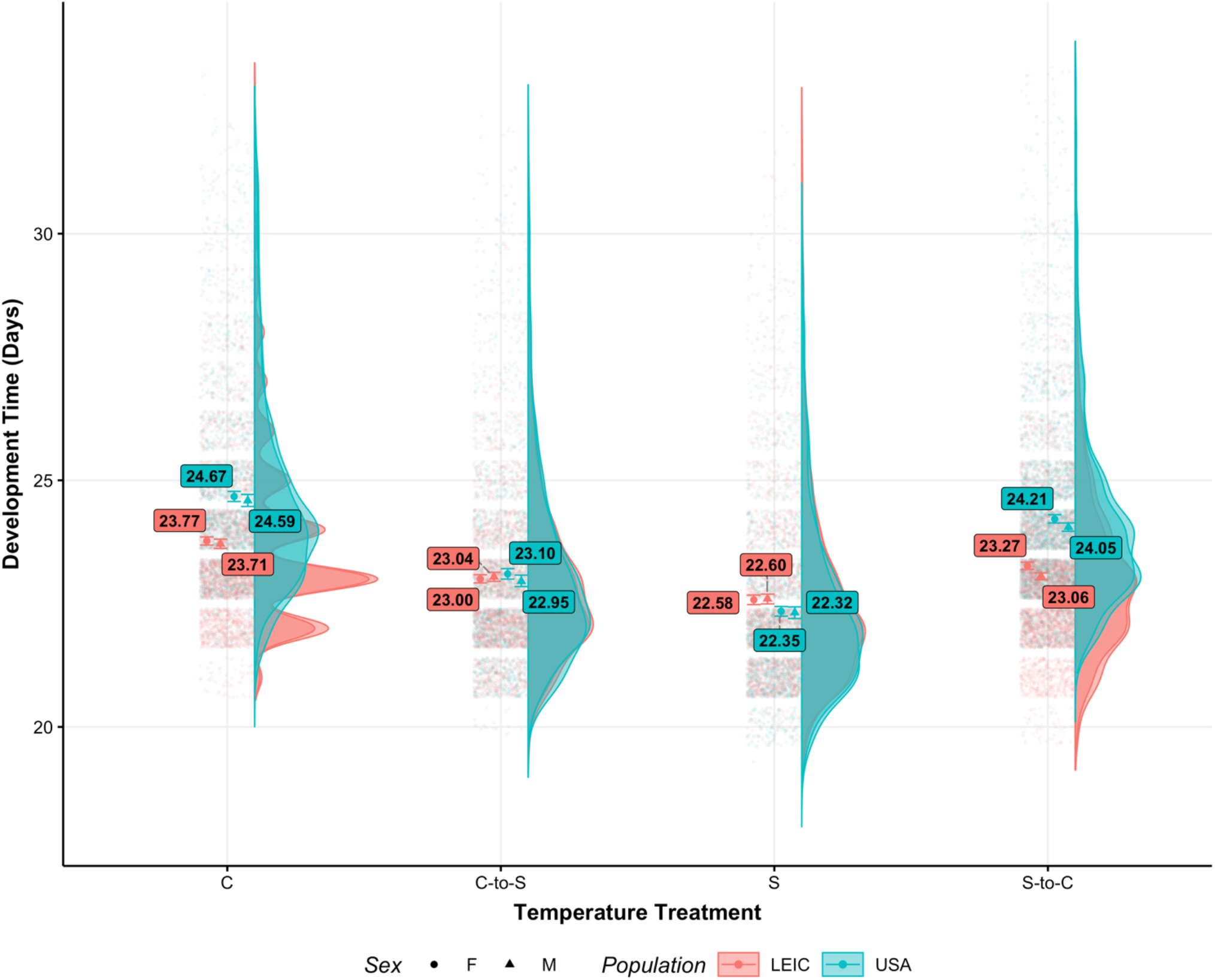
Development time in days of individuals from the Leicester (Red) or USA (Blue) populations in the C (Control), C-to-S (Control-to-Stress), S (Stress), or S-to-C (Stress-to-Control) temperature treatments. Sex is denoted with either circle (Female) or triangle (Male) points. Points represent mean values with accompanying standard errors. Half violins with associated jittered points show the distribution of raw data.

### Lifetime reproductive success

LRS was significantly influenced by the interaction between Population and Treatment (χ^2^ (1) = 12.61, *p* = 0.005). Across all temperature treatments, LRS was lower in the USA population than in the Leicester population (Average LRS: LEI = 53.2, USA = 40.2, Ratio = 1.32, *p*<0.001; Fig. 3A, Tables S3A-B). The largest differences between populations occurred in the C-to-S and C treatments (LEI-C-to-S = 50.2, USA-C-to-S = 33.6, Ratio = 1.49, *p*<0.001; LEI-C = 52.6, USA-C = 37.2, Ratio = 1.41, *p*<0.001; Fig. 3A, Tables S3A-B). Despite these differences, LRS was highest in the S-to-C treatment for both populations (LEI-C = 52.6, LEI-S-to-C = 69.7, Ratio = 0.75, *p*<0.001; USA-C = 37.2, USA-S-to-C = 60.9, Ratio = 0.61, *p*<0.001; Fig. 3A, Tables S3A-B).

**Figure 3.**
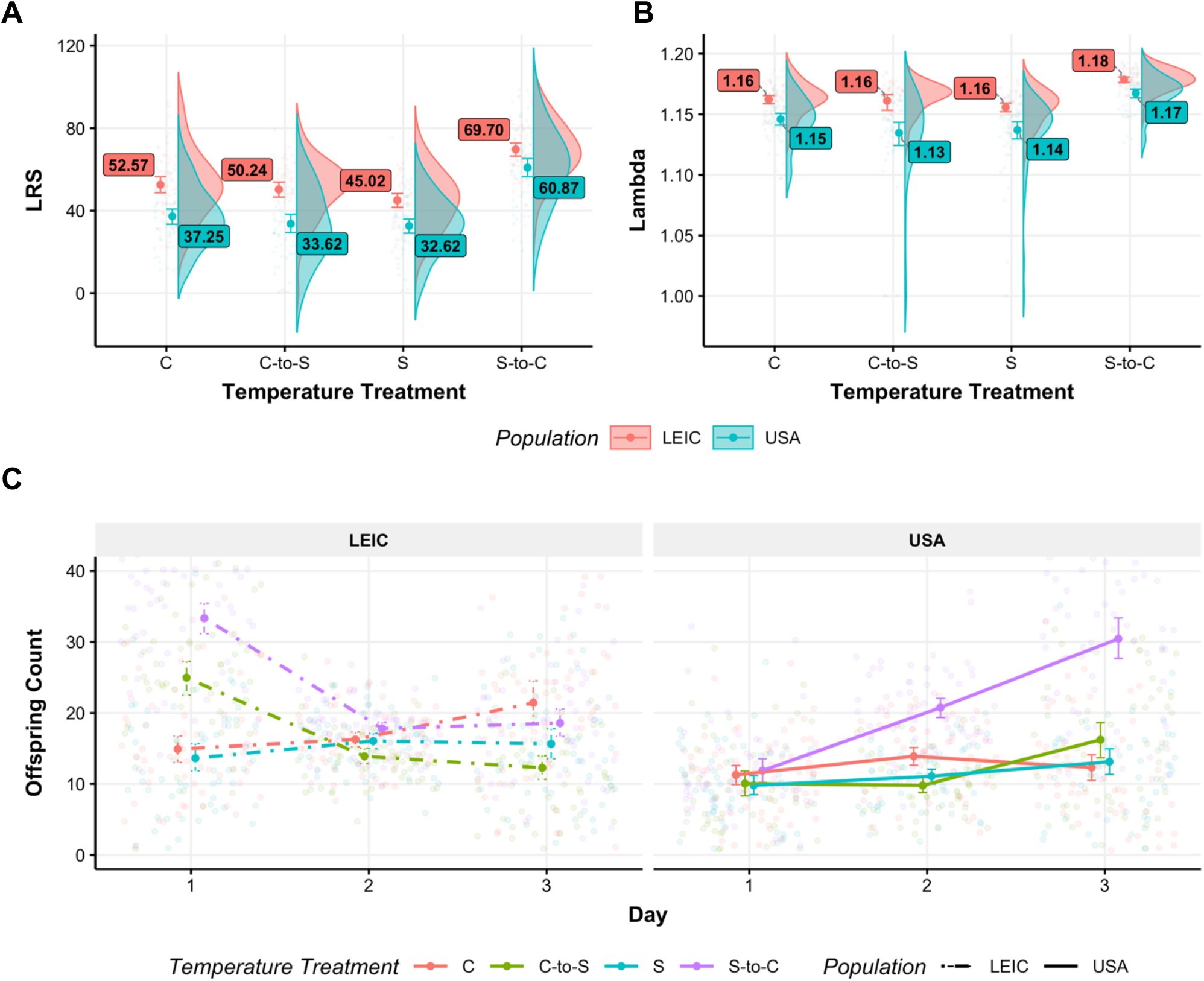
**(A)** Lifetime reproductive success (LRS) and **(B)** fitness (Lambda) of individuals from the Leicester (Red) or USA (Blue) populations in the C (Control), C-to-S (Control-to-Stress), S (Stress), or S-to-C (Stress-to-Control) temperature treatments. Points represent mean values with accompanying standard errors. Half violins show the distribution of raw data. **(C)** Age-specific reproduction of individuals from Leicester (dash-dot) or USA (solid) populations in the C (Control, Red), C-to-S (Control-to-Stress, Green), S (Stress, Blue), or S-to-C (Stress-to-Control, Purple) temperature treatments. Points with error bars represent mean values with accompanying standard errors. Jittered points represent raw data.

### Age-specific reproduction

The interaction between population, treatment and day significantly influenced age-specific reproduction (χ^2^ (3) = 203.54, *p*<0.001, Fig. 3C). Whilst continuous C and S treatments exhibited similar age-specific patterns across both populations (Fig. 3C), there were large differences between S-to-C and C-to-S treatments. This was particularly apparent on Days 1 and 3 of reproduction (Fig. 3C). Individuals from both LEI-S-to-C and LEI-C-to-S treatments exhibited far greater Day 1 reproduction than all other treatments from both populations (including the analogous S-to-C and C-to-S treatments). In contrast, USA-S-to-C exhibited increased reproduction *after* Day 1, with the result that Day 3 of reproduction was comparable with the Day 1 reproduction for LEI S-to-C (Fig. 3C). This variation in reproductive schedule resulted in large differences in age-specific slopes between the USA S-to-C and LEI S-to-C treatments (Average change in average offspring produced per day: LEI S-to-C = - 7.14, USA S-to-C = 8.78, estimate = −15.92, p<0.001; Fig. 3C, Tables S4A-B).

### Individual fitness

Whilst the individual main effects of Population and Temperature treatment significantly influenced lind (χ^2^ (1) = 81.02, *p* <0.001, χ^2^ (3) = 111.23, *p* <0.001, respectively; Fig. 3B), the interaction between these two did not (χ^2^ (3) = 7.64, *p* = 0.054). In a similar manner to LRS, the USA population had reduced lind over all temperature treatments (Average lind: LEI = 1.164, USA = 1.146, estimate = 0.018, p<0.001; Fig. 3B, Tables S5A-B). Across both populations, individuals from the S-to-C treatment had the highest lind (C = 1.154, C-to-S = 1.148, S = 1.146, S-to-C = 1.173, all comparisons p<0.001; Tables S5A-B). This was also consistent when looking within each population (all comparisons p<0.001).

## Discussion

The overall aim of this experiment was to evaluate the long- and short-term impacts of realistic stressful thermal fluctuations, conditions that are becoming biologically more and more relevant, on life history parameters of populations experiencing both constant (remaining within the same treatment group) and switching (changing to a different treatment group) thermal regimes. Previous hypotheses (Lewontin, 1974; Nevo, 1978) have suggested that evolution in fluctuating environments should select for genetically variable individuals that are robust to a wide range of thermal conditions with wide thermal performance curves (TPCs; Kassen, 2002; Jin & Agustí, 2018). In this experiment, individuals exposed to novel benign conditions (S-to-C) (although not those exposed to continuously stressful environments, S, see below) had higher reproductive fitness (higher LRS and lind) than those that were exposed to novel stressful conditions (C-to-S) or kept continuously in a benign environment (C). These results are broadly in line with previous research conducted in the opportunistic bacterial pathogen, *Serratia marcescens* (Ketola *et al.*, 2013). Heterogenous environments selected for bacterial clones that had faster growth and higher yield than those grown under constant environments, even when introduced into the same control temperature of 31 °C (Ketola *et al.*, 2013). This increased growth may be due to far greater selective pressure to remove mutations that could otherwise accumulate in more constant benign environments (Ketola *et al.*, 2013). However, results from a digital evolution experiment have suggested that this may only allow for optimization in short-term rather than long-term evolvability (Canino-Koning *et al.*, 2019).

Moreover our results suggest that individuals were able to tolerate and survive at wide thermal breadth, presumably owing to the evolution of phenotypic plasticity, but they reached maximal performance and experienced increased LRS only when conditions were optimal (Gilchrist, 1995; Ketola *et al.*, 2013). This could in part be due to differential gene expression as a result of exposure to a novel thermal environment (also known as cryptic genetic variability, McGuigan & Sgro, 2009; Paaby & Rockman, 2014). Similarly, this release of cryptic genetic variation could explain why individuals that were predicted to perform poorly when switching from a benign control to stressful fluctuating environment (C-to-S) suffered no fitness cost. However, the relative merits of this form of variation have been queried and require additional empirical testing (McGuigan & Sgro, 2009; Edwards & Yang, 2021; Johansson *et al.*, 2021). We note that not detecting costs of phenotypic plasticity does not mean that they do not exist. There might not be a sustainable cost to fitness if individuals are able to accurately predict future environmental conditions and avoid the expression of suboptimal genes (Auld *et al.*, 2010; Hoffmann & Bridle, 2022). Quantifying these costs would require an experimental design with additional thermal regimes beyond the scope of this current experiment.

When comparing body mass across treatments, individuals from the S-to-C group were significantly smaller (across both populations and sexes), despite having similar development times to those of the constant control group. Smaller body size could therefore be a cost of adaptation (Kassen, 2002; Ketola *et al.*, 2013) to the fluctuating conditions that is only detected in the benign environment and results in reduced body size (albeit with no negative effects on fitness). However, whether this reduction in mass is a disadvantage is difficult to determine. Large size provides a fitness advantage due to sexual selection (Andersson & Iwasa, 1996) and offers protection from desiccation (Le Lagadec *et al.*, 1998; Chown & Gaston, 2010) whereas small size decreases larval competition (Messina, 2004) and reduces heat stress (Chown & Gaston, 2010). Interestingly, similar to Hallsson & Björklund (2012a), we found that sexual dimorphism in adult body size decreased in the continuously stressful environment (S), although this could be considered population-specific as individuals from USA failed to show such a decrease.

Lastly, there were consistent population-level differences across all environmental treatments. Beetles from the USA population produced fewer offspring, took longer to develop (although this difference disappeared under stressful conditions), and had greater sexual dimorphism in body size compared to individuals from the Leicester population. Such genotypic variation between populations of the same species (particularly in *Callosobruchus maculatus*) has been found to impact larval competition (Messina, 1991), various morphometric measures (Rankin & Arnqvist, 2008), egg-to-adult survival and offspring production (Rankin & Arnqvist, 2008), and adult longevity and mortality (Fox *et al.*, 2004a; b). However, in this experiment, one particular observation stood out. Whilst a switch from a stressful environment to a benign regime (S-to-C) was associated with an increase in LRS and lind in both USA and Leicester populations, there was a noticeable difference in the age-specific reproductive schedules between the USA and Leicester populations. Leicester beetles produced more offspring on the first day and fewer on days two and three, whilst in the USA population the opposite was observed. Why such a difference in reproductive schedule exists deserves deeper investigation. If genetic variance differed between populations, this may explain why Leicester individuals were able to respond more quickly, with greater Day 1 reproduction. This could also in part explain why development time was similar across all experimental temperature regimes for the Leicester population. Together this clearly highlights the benefits of comparing and contrasting results from multiple strains, as the response of species to environmental change can often be highly population-specific.

As we have shown, successfully mimicking natural conditions within the laboratory is key to understanding how species will react to climate change. As temperatures increase and become more variable, the incorporation of more realistic conditions into laboratory studies such as these will allow us not only to predict more accurately how wild species will respond to climate change, specifically thermal stress, but also to produce accurate species- and population-specific thermal reaction norms (Paaijmans *et al.*, 2013; Buckley & Kingsolver, 2021). Therefore, this study further reinforces the importance of considering natural temperature conditions in order to better understand population- and species-specific responses to climate change both in response to short- and long-term environmental changes. Beyond this, there is also further need to extend thermal reaction norms to include additional environmental variables that may also vary in response to climate change (Buckley & Kingsolver, 2021).

## Acknowledgements

Funding from the American University of Paris supported this study. Many thanks to Sophie Bricout for helping to maintain the beetle populations and to Sophie Bricout, Linda Martz, Shannon Monahan, Sarah Sidi, and Xaviera Steele for their assistance with the experimental assays. Thanks also to Ram Vasudeva, Kynan Delaney, and Joel Pick for helpful discussions.

## Notes

### Competing Interest Statement

The authors have declared no competing interest.

### Summary of Updates

Manuscript has been updated with a reframed introduction and discussion.

## References

Andersson, M. & Iwasa, Y. 1996. Sexual selection. Trends Ecol. Evol. 11: 53–58. Elsevier.

Auld, J.R., Agrawal, A.A. & Relyea, R.A. 2010. Re-evaluating the costs and limits of adaptive phenotypic plasticity. Proc. R. Soc. B Biol. Sci. 277: 503–511.

Bauerfeind, S.S. & Fischer, K. 2014. Simulating climate change: temperature extremes but not means diminish performance in a widespread butterfly. Popul. Ecol. 56: 239–250.

Beck, C.W. & Blumer, L.S. 2014. A handbook on bean beetles, Callosobruchus maculatus. Natl. Sci. Found. URL Httpwww Beanbeetles Orghandbook Pdf Last Accessed 16 June 2015.

Berger, D., St\aangberg, J., Baur, J. & Walters, R.J. 2021. Elevated temperature increases genome-wide selection on de novo mutations. Proc. R. Soc. B 288: 20203094. The Royal Society.

Berger, D., Stångberg, J., Grieshop, K., Martinossi-Allibert, I. & Arnqvist, G. 2017. Temperature effects on life-history trade-offs, germline maintenance and mutation rate under simulated climate warming. Proc. R. Soc. B Biol. Sci. 284: 20171721. Royal Society.

Brakefield, P.M. & Mazzotta, V. 1995. Matching field and laboratory environments: effects of neglecting daily temperature variation on insect reaction norms. J. Evol. Biol. 8: 559–573.

Brooks, M.E., Kristensen, K., van Benthem, K.J., Magnusson, A., Berg, C.W., Nielsen, A., et al. 2017. glmmTMB balances speed and flexibility among packages for zero-inflated generalized linear mixed modeling. R J., doi: 10.32614/rj-2017-066.

Buckley, L.B. & Kingsolver, J.G. 2021. Evolution of Thermal Sensitivity in Changing and Variable Climates. Annu. Rev. Ecol. Evol. Syst. 52: 563–586.

Canino-Koning, R., Wiser, M.J. & Ofria, C. 2019. Fluctuating environments select for short-term phenotypic variation leading to long-term exploration. PLOS Comput. Biol. 15: e1006445. Public Library of Science.

Chang, M., Zhang, C., Li, M., Dong, J., Li, C., Liu, J., et al. 2022. Warming, temperature fluctuations and thermal evolution change the effects of microplastics at an environmentally relevant concentration. Environ. Pollut. 292:118363.

Chen, H., Zheng, X., Luo, M., Guo, J., Solangi, G.S., Wan, F., et al. 2018. Effect of short-term high-temperature exposure on the life history parameters of Ophraella communa.

Chown, S.L. & Gaston, K.J. 2010. Body size variation in insects: a macroecological perspective. Biol. Rev. 85: 139–169.

Colinet, H., Sinclair, B.J., Vernon, P. & Renault, D. 2015. Insects in Fluctuating Thermal Environments. Annu. Rev. Entomol. 60: 123–140.

Deutsch, C.A., Tewksbury, J.J., Huey, R.B., Sheldon, K.S., Ghalambor, C.K., Haak, D.C., et al. 2008. Impacts of climate warming on terrestrial ectotherms across latitude. Proc. Natl. Acad. Sci. 105: 6668–6672.

Edwards, C.B. & Yang, L.H. 2021. Evolved Phenological Cueing Strategies Show Variable Responses to Climate Change. Am. Nat. 197: E1–E16. The University of Chicago Press.

Fischer, K., Kölzow, N., Höltje, H. & Karl, I. 2011. Assay conditions in laboratory experiments: is the use of constant rather than fluctuating temperatures justified when investigating temperature-induced plasticity? Oecologia 166: 23–33.

Folguera, G., Bastías, D.A., Caers, J., Rojas, J.M., Piulachs, M.-D., Bellés, X., et al. 2011. An experimental test of the role of environmental temperature variability on ectotherm molecular, physiological and life-history traits: implications for global warming. Comp. Biochem. Physiol. A. Mol. Integr. Physiol. 159: 242–246.

Fox, C.W. 1993. Multiple mating, lifetime fecundity and female mortality of the bruchid beetle, Callosobruchus maculatus (Coleoptera: Bruchidae). Funct. Ecol. 203–208. JSTOR.

Fox, C.W., Bush, M.L., Roff, D.A. & Wallin, W.G. 2004a. Evolutionary genetics of lifespan and mortality rates in two populations of the seed beetle, Callosobruchus maculatus. Heredity 92: 170–181.

Fox, C.W., Czesak, M.E. & Wallin, W.G. 2004b. Complex genetic architecture of population differences in adult lifespan of a beetle: Nonadditive inheritance, gender differences, body size and a large maternal effect. J. Evol. Biol. 17: 1007–1017.

Fox, J., Weisberg, S., Adler, D., Bates, D., Baud-Bovy, G., Ellison, S., et al. 2012. Package ‘car.’ Vienna R Found. Stat. Comput.

Fricke, C. & Arnqvist, G. 2004. Divergence in replicated phylogenies: the evolution of partial post-mating prezygotic isolation in bean weevils. J. Evol. Biol. 17: 1345–1354. Wiley Online Library.

Gilchrist, G.W. 1995. Specialists and Generalists in Changing Environments. I. Fitness Landscapes of Thermal Sensitivity. Am. Nat. 146: 252–270.

Hallsson, L.R. & Björklund, M. 2012. Selection in a fluctuating environment and the evolution of sexual dimorphism in the seed beetle Callosobruchus maculatus. J. Evol. Biol. 25: 1564–1575.

Hartig, F. 2020. DHARMa: Residual Diagnostics for Hierarchical (Multi-Level / Mixed) Regression Models. R package.

Hetem, R.S., Fuller, A., Maloney, S.K. & Mitchell, D. 2014. Responses of large mammals to climate change. Temperature 1: 115–127.

Holt, R.D. 1990. The microevolutionary consequences of climate change. Trends Ecol. Evol. 5: 311–315. Elsevier.

Ivimey-Cook, E., Bricout, S., Candela, V., Maklakov, A.A. & Berg, E.C. 2021. Inbreeding reduces fitness of seed beetles under thermal stress. J. Evol. Biol. jeb.13899.

Janzen, D.H. 1967. Why Mountain Passes are Higher in the Tropics. Am. Nat. 101: 233–249. [University of Chicago Press, American Society of Naturalists].

Jin, P. & Agustí, S. 2018. Fast adaptation of tropical diatoms to increased warming with trade-offs. Sci. Rep. 8: 17771.

Johansson, F., Watts, P.C., Sniegula, S. & Berger, D. 2021. Natural selection mediated by seasonal time constraints increases the alignment between evolvability and developmental plasticity. Evolution 75: 464–475.

Kassen, R. 2002. The experimental evolution of specialists, generalists, and the maintenance of diversity. J. Evol. Biol. 15: 173–190.

Kellermann, V., Chown, S.L., Schou, M.F., Aitkenhead, I., Janion-Scheepers, C., Clemson, A., et al. 2019. Comparing thermal performance curves across traits: how consistent are they? J. Exp. Biol. jeb.193433.

Kellermann, V. & van Heerwaarden, B. 2019. Terrestrial insects and climate change: adaptive responses in key traits. Physiol. Entomol. 44: 99–115.

Ketola, T., Mikonranta, L., Zhang, J., Saarinen, K., Örmälä, A.-M., Friman, V.-P., et al. 2013. FLUCTUATING TEMPERATURE LEADS TO EVOLUTION OF THERMAL GENERALISM AND PREADAPTATION TO NOVEL ENVIRONMENTS. Evolution 67: 2936–2944. [Society for the Study of Evolution, Wiley].

Klepsatel, P., Girish, T.N., Dircksen, H. & Ga, M. 2019. Reproductive fitness of Drosophila is maximised by optimal developmental temperature. J. Exp. Biol. 11.

Lashof, D.A. & Ahuja, D.R. 1990. Relative contributions of greenhouse gas emissions to global warming. Nature 344: 529–531. Springer.

Le Lagadec, M.D., Chown, S.L. & Scholtz, C.H. 1998. Desiccation resistance and water balance in southern African keratin beetles (Coleoptera, Trogidae): the influence of body size and habitat. J. Comp. Physiol. B 168: 112–122.

Lehmann, P., Ammunét, T., Barton, M., Battisti, A., Eigenbrode, S.D., Jepsen, J.U., et al. 2020. Complex responses of global insect pests to climate warming. Front. Ecol. Environ. 18: 141–150.

Lenth, R., Singmann, H., Love, J., Buerkner, P. & Herve, M. 2019. emmeans: Estimated Marginal Means, aka Least-Squares Means (Version 1.3. 4).

Leslie, P.H. 1945. On the Use of Matrices in Certain Population Mathematics. Biometrika 33: 183–212.

Lewontin, R.C. 1974. The genetic basis of evolutionary change. Columbia University Press New York.

Ma, C.-S., Ma, G. & Pincebourde, S. 2021a. Survive a Warming Climate: Insect Responses to Extreme High Temperatures. Annu. Rev. Entomol. 66: 163–184.

Ma, G., Hoffmann, A.A. & Ma, C.-S. 2021b. Are extreme high temperatures at low or high latitudes more likely to inhibit the population growth of a globally distributed aphid? J. Therm. Biol. 98: 102936.

MacLean, H.J., Sørensen, J.G., Kristensen, T.N., Loeschcke, V., Beedholm, K., Kellermann, V., et al. 2019. Evolution and plasticity of thermal performance: an analysis of variation in thermal tolerance and fitness in 22 Drosophila species. Philos. Trans. R. Soc. B 374: 20180548. The Royal Society.

Magnusson, A., Skaug, H.J., Nielsen, A., Berg, C., Kristensen, K., Maechler, M., et al. 2019. Package ‘glmmTMB. R Package Version 01.

Masson-Delmotte, V., Zhai, P. & Pirani, A. 2021. IPCC, 2021: Climate Change 2021: The Physical Science Basis. Contribution of Working Group I to the Sixth Assessment Report of the Intergovernmental Panel on Climate Change.

McGuigan, K. & Sgro, C.M. 2009. Evolutionary consequences of cryptic genetic variation. Trends Ecol. Evol. 24: 305–311. Elsevier.

Messina, F.J. 1991. Competitive interactions between larvae from divergent strains of the cowpea weevil (Coleoptera: Bruchidae). Environ. Entomol. 20: 1438–1443. Oxford University Press Oxford, UK.

Messina, F.J. 2004. Predictable Modification of Body Size and Competitive Ability Following a Host Shift by a Seed Beetle. Evolution 58: 2788–2797. [Society for the Study of Evolution, Wiley].

Mitchell, R. 1991. The traits of a biotype of Callosobruchus maculatus (F.)(Coleoptera: Bruchidae) from South India. J. Stored Prod. Res. 27: 221–224. Elsevier.

Morash, A.J., Neufeld, C., MacCormack, T.J. & Currie, S. 2018. The importance of incorporating natural thermal variation when evaluating physiological performance in wild species. J. Exp. Biol. 221: jeb164673.

Morgan Fleming, J., Carter, A.W. & Sheldon, K.S. 2021. Dung beetles show metabolic plasticity as pupae and smaller adult body size in response to increased temperature mean and variance. J. Insect Physiol. 131: 104215.

Nevo, E. 1978. Genetic variation in natural populations: Patterns and theory. Theor. Popul. Biol. 13: 121–177.

Niehaus, A.C., Angilletta, M.J., Sears, M.W., Franklin, C.E. & Wilson, R.S. 2012. Predicting the physiological performance of ectotherms in fluctuating thermal environments. J. Exp. Biol. 215: 694–701.

Paaby, A.B. & Rockman, M.V. 2014. Cryptic genetic variation: evolution’s hidden substrate. Nat. Rev. Genet. 15: 247–258.

Paaijmans, K.P., Heinig, R.L., Seliga, R.A., Blanford, J.I., Blanford, S., Murdock, C.C., et al. 2013. Temperature variation makes ectotherms more sensitive to climate change. Glob. Change Biol. 19: 2373–2380.

Pachauri, R.K., Allen, M.R., Barros, V.R., Broome, J., Cramer, W., Christ, R., et al. 2014. Climate change 2014: synthesis report. Contribution of Working Groups I, II and III to the fifth assessment report of the Intergovernmental Panel on Climate Change. Ipcc.

Parmesan, C. 2006. Ecological and Evolutionary Responses to Recent Climate Change. Annu. Rev. Ecol. Evol. Syst. 37: 637–669.

Pereira, H.M., Navarro, L.M. & Martins, I.S. 2012. Global Biodiversity Change: The Bad, the Good, and the Unknown. Annu. Rev. Environ. Resour. 37: 25–50.

R Core Team. 2016. R: A language and environment for statistical computing. R Found. Stat. Comput. Vienna Austria 0: {ISBN} 3-900051-07-0.

Rankin, D.J. & Arnqvist, G. 2008. Sexual Dimorphism Is Associated with Population Fitness in the Seed Beetle Callosobruchus Maculatus. Evolution 62: 622–630.

Rogell, B., Widegren, W., Hallsson, L.R., Berger, D., Björklund, M. & Maklakov, A.A. 2014. Sex-dependent evolution of life-history traits following adaptation to climate warming. Funct. Ecol. 28: 469–478.

Stubben, C. & Milligan, B. 2007. Estimating and analyzing demographic models using the popbio package in R. J. Stat. Softw. 22: 1–23.

Thompson, R.M., Beardall, J., Beringer, J., Grace, M. & Sardina, P. 2013. Means and extremes: building variability into community-level climate change experiments. Ecol. Lett. 16: 799–806.

Trisos, C.H., Merow, C. & Pigot, A.L. 2020. The projected timing of abrupt ecological disruption from climate change. Nature 580: 496–501. Nature Publishing Group.

Ummenhofer, C.C. & Meehl, G.A. 2017. Extreme weather and climate events with ecological relevance: a review. Philos. Trans. R. Soc. B Biol. Sci. 372: 20160135. Royal Society.

Ursprung, C., Den Hollander, M. & Gwynne, D.T. 2009. Female seed beetles, Callosobruchus maculatus, remate for male-supplied water rather than ejaculate nutrition. Behav. Ecol. Sociobiol. 63: 781–788. Springer.

Vasseur, D.A., DeLong, J.P., Gilbert, B., Greig, H.S., Harley, C.D.G., McCann, K.S., et al. 2014. Increased temperature variation poses a greater risk to species than climate warming. Proc. R. Soc. B Biol. Sci. 281: 20132612.

Vasudeva, R., Deeming, D.C. & Eady, P.E. 2014. Developmental temperature affects the expression of ejaculatory traits and the outcome of sperm competition in Callosobruchus maculatus. J. Evol. Biol. 27: 1811–1818.

Visser, M.E., Noordwijk, A.J. van, Tinbergen, J.M. & Lessells, C.M. 1998. Warmer springs lead to mistimed reproduction in great tits (Parus major). Proc. R. Soc. Lond. B Biol. Sci. 265: 1867–1870. Royal Society.

Visser, M.E., te Marvelde, L. & Lof, M.E. 2012. Adaptive phenological mismatches of birds and their food in a warming world. J. Ornithol. 153: 75–84.

Walsh, B.S., Parratt, S.R., Hoffmann, A.A., Atkinson, D., Snook, R.R., Bretman, A., et al. 2019. The Impact of Climate Change on Fertility. Trends Ecol. Evol. 34: 249–259.

Walters, R.J., Blanckenhorn, W.U. & Berger, D. 2012. Forecasting extinction risk of ectotherms under climate warming: an evolutionary perspective. Funct. Ecol. 26: 1324–1338.

Wickham, H. 2009. ggplot2 Elegant Graphics for Data Analysis.

