## Supplementary Material for "Genetic and thermal variation influence adaptation to fluctuating temperature in the seed beetle, *Callosobruchus maculatus*"

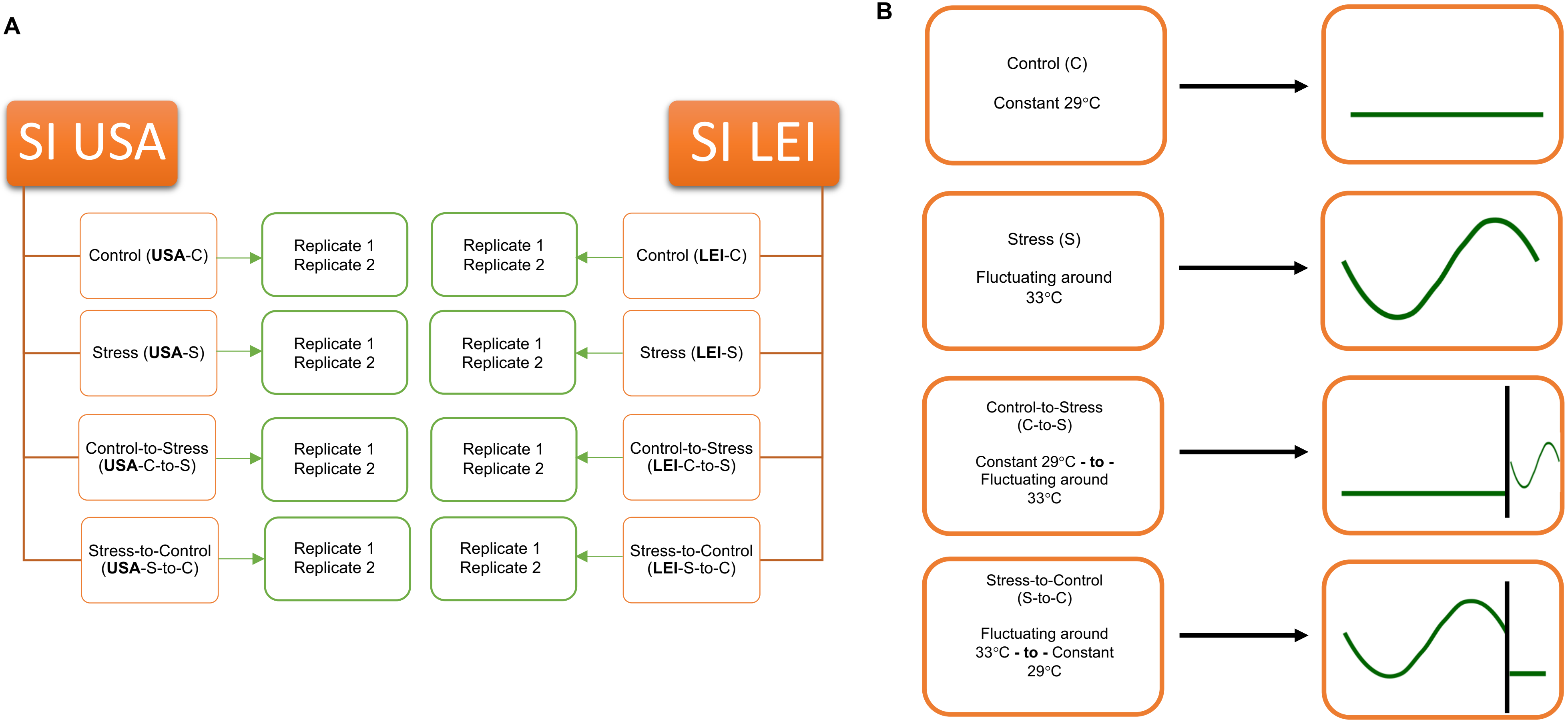


**Figure S1. (A)** A schematic diagram of the four temperature treatments (Control, Stress, Control-to-Stress, and Stress-to-Control) across both the USA (SI USA) and Leicester (SI LEI) populations. **(B)** A representation of the four different temperature treatments used. The black bar in the bottom two treatments denotes a switch from one treatment to another.

**Table S1A.** Full model output from the mixed effect model for body mass.

| ***Predictors*** | ***Estimates*** | ***std. Error*** | ***Statistic*** | ***p*** |
|---|---|---|---|---|
| (Intercept) | 0.09 | 0.00 | 52.99 | **<0.001** |
| Population [USA] | 0.00 | 0.00 | 2.32 | **0.020** |
| Treatment [C-to-S] | 0.00 | 0.00 | 1.69 | 0.091 |
| Treatment [S] | 0.00 | 0.00 | 0.63 | 0.531 |
| Treatment [S-to-C] | -0.01 | 0.00 | -2.41 | **0.016** |
| Sex [M] | -0.01 | 0.00 | -7.61 | **<0.001** |
| Replicate [2] | 0.00 | 0.00 | 3.04 | **0.002** |
| Population [USA] * Treatment [C-to-S] | -0.00 | 0.00 | -1.51 | 0.132 |
| Population [USA] * Treatment [S] | -0.00 | 0.00 | -1.03 | 0.305 |
| Population [USA] * Treatment [S-to-C] | -0.00 | 0.00 | -0.98 | 0.330 |
| Population [USA] * Sex [M] | -0.01 | 0.00 | -5.09 | **<0.001** |
| Treatment [C-to-S] * Sex [M] | -0.00 | 0.00 | -0.56 | 0.578 |
| Treatment [S] * Sex [M] | 0.01 | 0.00 | 3.51 | **<0.001** |
| Treatment [S-to-C] * Sex [M] | -0.00 | 0.00 | -0.41 | 0.679 |
| **Observations** | 144 | | | |

**Table S1B.** Pairwise comparisons of relevant treatment contrasts using emmeans.

| *Contrast* | *Estimate* | *std. Error* | *df* | *p* |
|---|---|---|---|---|
| *F-LEI vs M-LEI* | 0.0126 | 0.0013 | 129 | **<0.001** |
| *F-USA vs M-USA* | 0.0213 | 0.0012 | 129 | **<0.001** |
| *F-LEI vs F-USA* | -0.0020 | 0.0012 | 129 | 0.347 |
| *M-LEI vs M-USA* | 0.0067 | 0.0012 | 129 | **<0.001** |
| *F-S-to-C vs F-C* | -0.0063 | 0.0017 | 129 | **0.006** |
| *F-S-to-C vs F-S* | -0.0064 | 0.0018 | 129 | **0.011** |
| *F-S-to-C vs F-C-to-S* | -0.0081 | 0.0018 | 129 | **<0.001** |
| *M-S-to-C vs M-C* | -0.0072 | 0.0017 | 129 | **<0.001** |
| *M-S-to-C vs M-S* | -0.0155 | 0.0018 | 129 | **<0.001** |
| *M-S-to-C vs M-C-to-S* | -0.0077 | 0.0018 | 129 | **<0.001** |
| *F-C vs M-C* | 0.0185 | 0.0015 | 129 | **<0.001** |
| *F-S-to-C vs M-S-to-C* | 0.0194 | 0.0018 | 129 | **<0.001** |
| *F-S vs M-S* | 0.0103 | 0.0018 | 129 | **<0.001** |
| *F-C-to-S vs M-C-to-S* | 0.0198 | 0.0018 | 129 | **<0.001** |

**Table S2A.** Full model output from the mixed effect model for development time.

| *Predictors* | *Estimates* | *std. Error* | *Statistic* | *p* |
|---|---|---|---|---|
| (Intercept) | 3.16 | 0.00 | 939.18 | **<0.001** |

| | | | | |
|---|---|---|---|---|
| Population [USA] | 0.04 | 0.00 | 9.34 | **<0.001** |
| Treatment [C-to-S] | -0.03 | 0.00 | -6.95 | **<0.001** |
| Treatment [S] | -0.05 | 0.00 | -11.09 | **<0.001** |
| Treatment [S-to-C] | -0.02 | 0.00 | -4.69 | **<0.001** |
| Sex [M] | -0.00 | 0.00 | -1.52 | 0.128 |
| Replicate [2] | 0.00 | 0.00 | 1.01 | 0.314 |
| Population [USA] * Treatment [C-to-S] | -0.04 | 0.01 | -6.37 | **<0.001** |
| Population [USA] * Treatment [S] | -0.05 | 0.01 | -8.07 | **<0.001** |
| Population [USA] * Treatment [S-to-C] | -0.00 | 0.01 | -0.23 | 0.814 |
| Population [USA] * Sex [M] | -0.00 | 0.00 | -0.85 | 0.395 |
| Treatment [C-to-S] * Sex [M] | 0.00 | 0.00 | 0.79 | 0.428 |
| Treatment [S] * Sex [M] | 0.00 | 0.00 | 1.33 | 0.182 |
| Treatment [S-to-C] * Sex [M] | -0.00 | 0.00 | -1.73 | 0.084 |
| **Random Effects** | | | | |
| **$\sigma^2$** | 0.00 | | | |
| **$\tau_{00}$ ID** | 0.00 | | | |
| **N ID** | 474 | | | |
| **Observations** | 22636 | | | |

**Table S2B.** Pairwise comparisons of relevant treatment contrasts using emmeans.

| ***Contrast*** | ***Estimate*** | ***std. Error*** | ***df*** | ***p*** |
|---|---|---|---|---|
| *LEI-C vs USA-C* | 0.960 | 0.0042 | 22620 | **<0.001** |
| *LEI-S-to-C vs USA- S-to-C* | 0.961 | 0.0040 | 22620 | **<0.001** |
| *LEI-S vs USA-S* | 1.010 | 0.0045 | 22620 | 0.378 |
| *LEI-C-to-S vs USA-C-to-S* | 0.999 | 0.0045 | 22620 | 1.000 |
| *F-S-to-C vs M-S-to-C* | 1.008 | 0.0016 | 225620 | **<0.001** |

**Table S3A.** Full model output from the mixed effect model for lifetime reproductive success (LRS)

No zero-inflation detected (p = 1). Conway-Maxwell Poisson identified as best model.

| ***Predictors*** | ***Estimates*** | ***std. Error*** | ***Statistic*** | ***p*** |
|---|---|---|---|---|
| (Intercept) | 3.94 | 0.04 | 91.13 | **<0.001** |
| Population [USA] | -0.34 | 0.06 | -5.51 | **<0.001** |
| Treatment [C-to-S] | -0.05 | 0.06 | -0.78 | 0.437 |
| Treatment [S] | -0.16 | 0.06 | -2.58 | **0.010** |
| Treatment [S-to-C] | 0.28 | 0.05 | 5.25 | **<0.001** |
| Replicate [2] | 0.04 | 0.03 | 1.20 | 0.232 |
| Population [USA] * Treatment [C-to-S] | -0.06 | 0.09 | -0.62 | 0.534 |
| Population [USA] * Treatment [S] | 0.02 | 0.09 | 0.24 | 0.811 |

| | | | | |
|---|---|---|---|---|
| Population [USA] * Treatment [S-to-C] | 0.21 | 0.08 | 2.57 | **0.010** |
| **Observations** | 474 | | | |

**Table S3B.** Pairwise comparisons of relevant treatment contrasts using emmeans.

| ***Contrast*** | ***Estimate*** | ***std. Error*** | ***df*** | ***p*** |
|---|---|---|---|---|
| *LEI vs USA* | 1.320 | 0.0408 | 467 | **<0.001** |
| *LEI-C-to-S vs USA-C-to-S* | 1.493 | 0.0985 | 464 | **<0.001** |
| *LEI-C vs USA-C* | 1.411 | 0.0882 | 464 | **<0.001** |
| *LEI-C vs LEI-S-to-C* | 0.754 | 0.0406 | 464 | **<0.001** |
| *USAI-C vs USA-S-to-C* | 0.612 | 0.0372 | 464 | **<0.001** |

**Table S4A.** Full model output from the mixed effect model for age-specific reproduction.

No zero-inflation detected (p = 1). Conway-Maxwell Poisson identified as best model.

| ***Predictors*** | ***Estimates*** | ***std. Error*** | ***Statistic*** | ***p*** |
|---|---|---|---|---|
| (Intercept) | 2.45 | 0.09 | 28.64 | **<0.001** |
| Population [USA] | -0.04 | 0.13 | -0.31 | 0.757 |
| Treatment [C-to-S] | 1.08 | 0.11 | 9.38 | **<0.001** |
| Treatment [S] | 0.10 | 0.12 | 0.81 | 0.417 |
| Treatment [S-to-C] | 1.28 | 0.11 | 11.86 | **<0.001** |

| | | | | |
|---|---|---|---|---|
| Day | 0.19 | 0.04 | 5.16 | **<0.001** |
| Replicate [2] | 0.02 | 0.03 | 0.89 | 0.372 |
| Population [USA] * Treatment [C-to-S] | -1.58 | 0.18 | -8.60 | **<0.001** |
| Population [USA] * Treatment [S] | -0.41 | 0.19 | -2.23 | **0.026** |
| Population [USA] * Treatment [S-to-C] | -1.64 | 0.17 | -9.74 | **<0.001** |
| Population [USA] * Day | -0.15 | 0.06 | -2.63 | **0.009** |
| Treatment [C-to-S] * Day | -0.57 | 0.05 | -10.85 | **<0.001** |
| Treatment [S] * Day | -0.12 | 0.05 | -2.23 | **0.025** |
| Treatment [S-to-C] * Day | -0.51 | 0.05 | -10.56 | **<0.001** |
| (Population [USA] * Treatment [C-to-S]) * Day | 0.79 | 0.08 | 9.73 | **<0.001** |
| (Population [USA] * Treatment [S]) * Day | 0.22 | 0.08 | 2.75 | **0.006** |
| (Population [USA] * Treatment [S-to-C]) * Day | 0.92 | 0.07 | 12.56 | **<0.001** |
| **Random Effects** | | | | |
| **$\sigma^2$** | 2.85 | | | |
| **$\tau_{00\ ID}$** | 0.03 | | | |
| **$N_{ID}$** | 474 | | | |
| **Observations** | 1388 | | | |

**Table S4B.** Pairwise comparisons of relevant treatment contrasts using emmeans.

| *Contrast* | *Estimate* | *std. Error* | *df* | *p* |
|---|---|---|---|---|
| *LEI-S-to-C vs USA-S-to-C* | -15.92 | 0.979 | 1369 | **<0.001** |

**Table S5A.** Full model output from the mixed effect model for individual fitness λind.

| *Predictors* | *Estimates* | *std. Error* | *Statistic* | *p* |
|---|---|---|---|---|
| (Intercept) | 1.16 | 0.00 | 473.00 | **<0.001** |
| Population [USA] | -0.02 | 0.00 | -9.00 | **<0.001** |
| Treatment [C-to-S] | -0.01 | 0.00 | -2.14 | **0.032** |
| Treatment [S] | -0.01 | 0.00 | -2.71 | **0.007** |
| Treatment [S-to-C] | 0.02 | 0.00 | 6.69 | **<0.001** |
| Replicate [2] | 0.00 | 0.00 | 1.20 | 0.232 |
| **Observations** | 474 | | | |

**Table S5B.** Pairwise comparisons of relevant treatment contrasts using emmeans.

| *Contrast* | *Estimate* | *std. Error* | *df* | *p* |
|---|---|---|---|---|
| *LEI vs USA* | 0.0181 | 0.0020 | 467 | **<0.001** |
| *S-to-C vs C* | 0.0183 | 0.0028 | 467 | **<0.001** |
| *S-to-C vs C-to-S* | 0.0250 | 0.0029 | 467 | **<0.001** |

| | | | | |
|---|---|---|---|---|
| *S-to-C vs C-to-S* | 0.0266 | 0.0029 | 467 | **<0.001** |
